# Mast cells impair melanoma cell homing and metastasis by inhibiting HMGA1 secretion

**DOI:** 10.1101/2022.05.31.494113

**Authors:** Alberto Benito-Martin, Marta Hergueta-Redondo, Laura Nogués, Elena Castellano-Sanz, Eduardo Garvin, Michele Cioffi, Paloma Sola-Castrillo, Weston Buehring, Pilar Ximénez-Embún, Javier Muñoz, Irina Matei, Josep Villanueva, Héctor Peinado

## Abstract

Metastatic disease is the major cause of death from cancer. From the primary tumor, cells remotely prepare the environment of the future metastatic sites by secreted factors and extracellular vesicles. During this process, known as pre-metastatic niche formation, immune cells play a crucial role. Mast cells are hematopoietic bone marrow-derived innate immune cells whose function in lung immune response to invading tumors remains to be defined. We found reduced melanoma lung metastasis in mast cell deficient mouse models (Wsh and MCTP5-Cre-RDTR), supporting a pro-metastatic role for mast cells *in vivo*. However, due to evidence pointing to their antitumorigenic role, we studied the impact of mast cells in melanoma cell function *in vitro*. Surprisingly, *in vitro* co-culture of bone-marrow derived mast cells with melanoma cells showed that they have an intrinsic anti-metastatic activity. Mass spectrometry analysis of melanoma-mast cell co-cultures secretome showed that HMGA1 secretion by melanoma cells was significantly impaired. Consistently, HMGA1 knock down in B16-F10 cells reduced their metastatic capacity *in vivo*. Importantly, analysis of HMGA1 expression in human melanoma tumors showed that metastatic tumors with high HMGA1 expression are associated with reduced overall and disease-free survival. Moreover, we show that HMGA1 is reduced in the nuclei and enriched in the cytoplasm of melanoma metastatic lesions when compared to primary tumors. These data suggest that high HMGA1 expression and secretion from melanoma cells promotes metastatic behavior. Targeting HMGA1 expression intrinsically or extrinsically by mast cells actions reduce melanoma metastasis. Our results pave the way to the use of HMGA1 as anti-metastatic target in melanoma as previously suggested in other cancer types.

## Introduction

Malignant melanoma (MM) is one of the most devastating cancers, and metastatic disease responsible for 90% of cancer related deaths(1, 2). Tumors induce the formation of pre-metastatic niches (PMN), specialized microenvironments in distant organs that are conducive to the survival and outgrowth of tumor cells (3). PMN formation is the result of the systemic effect of tumor-secreted factors and extracellular vesicles (EVs) that facilitate the survival and outgrowth of tumor cells in the target organ, collaborating with bone marrow-derived cells recruited to the metastatic site (3). Mast cells (MCs) are hematopoietic bone marrow-derived immune cells. The function in the host immune response to tumors remains to be defined (4, 5). Mast cells (MC) are broadly characterized by their content, rich in granules containing enzymes, cytokines, and proteases like histamine or tryptase (6, 7). MCs are found in most tissues, flourishing in barrier tissues, such skin, gut or lungs (8), playing protective roles(7), although they are associated with allergy and anaphylaxis (9). The role of MCs in tumorigenesis is controversial, as several studies have proposed that MCs contribute to tumor progression or metastasis (10-12), while others report MCs exerting anti-primary tumor effects (7, 13-16).

To interpret the dual role of MC in melanoma progression, we analyzed MC function in two scenarios: 1) *in vivo* analyzing tumor growth and metastasis using MC-deficient models and 2) *in vitro* exposing melanoma cells to bone marrow derived MCs (BMDMCs). On one hand we found that, in tumor-bearing mice, MCs have a pro-metastatic activity as denoted by metastasis reduction in mast cell-deficient models. Interestingly, on the other hand, BMDMCs from wild type (*wt*)mice have an intrinsic anti-metastatic effect when co-cultured with melanoma cells These data support a dual role for MCs during tumor progression: an innate anti-tumoral role reducing metastatic behavior, but once tumor progresses mast cells support metastatic spread in melanoma models. We focused on analyzing the potential MCs anti-metastatic role to develop novel therapeutic strategies. Mechanistically, MCs co-culture with melanoma cells inhibits HMGA1 secretion. HMGA1 is a nuclear protein that binds to the minor groove of AT-rich DNA strands and controls transcriptional activity of several genes(17). Previous reports showed that increased cytoplasm expression and secretion of HMGA1 is related to metastatic behavior in triple negative breast cancer (18). Ablation of HMGA1 expression in B16-F10 cells decreased melanoma lung metastasis *in vivo*. Analysis of HMGA1 expression in human melanoma primary tumors and metastasis showed that high levels of HMGA1 expression in metastases are associated with reduced overall and disease-free survival. Finally, HMGA1 expression analysis in a cohort of 36 tumors and 12 metastatic lesions showed that cytoplasmic staining was enriched in metastatic lesions compared to primary tumors, suggesting that increased levels and cytoplasmic localization of HMGA1 are associated to metastatic behavior in melanoma. Overall, our data support a model where HMGA1 enhances metastatic ability along melanoma progression being mobilized from nuclei to cytoplasm and secreted in melanoma cells. MCs interaction with tumor cells or HMGA1 interference reduced metastatic spread in melanoma cells suggesting that HMGA1 targeting strategies may represent a novel mechanism to reduce metastatic behavior in melanoma.

## Results

### Mast cells reinforce melanoma metastasis in tumor-bearing mice

To determine the role of mast cells (MCs) in melanoma metastasis we subcutaneously injected (sc.i) lung metastatic B16-F10-mCherry cells in mast cell deficient mice (Wsh). MC-deficient mice developed bigger tumors than *wt* mice (Figure 1A). Analysis of spontaneous metastasis showed that lung metastasis was reduced in MCs-deficient mice regardless of primary tumor growth (Figure 1B). We quantified mCherry expression by qRT-PCR observing a significant reduction of metastatic tumor cells (Fig. 1C). Since Wsh mice have defects in hematopoietic progenitors in addition to MC we used an additional mouse model, the MCTP5-Cre-RDTR mouse model, deficient in connective-tissue MCs (19). This model has been previously used to demonstrate the reduction of experimental melanoma metastasis (16), but not in spontaneous metastatic models. We first verified decreased MC numbers in lungs (Fig 1D) of MCTP5-Cre-RDTR mice. We next analyzed primary tumor growth and spontaneous metastasis after B16F10-mCherry sc.i. in MCTP5-Cre-RDTR (Fig. 1E, 1F). We observed that while mice did not present primary tumor growth differences (Fig 1E), there was a significant reduction in spontaneous lung metastasis (Fig. 1F). These data support a pro-metastatic role for MCs in tumor-bearing mice.

**Figure 1.**
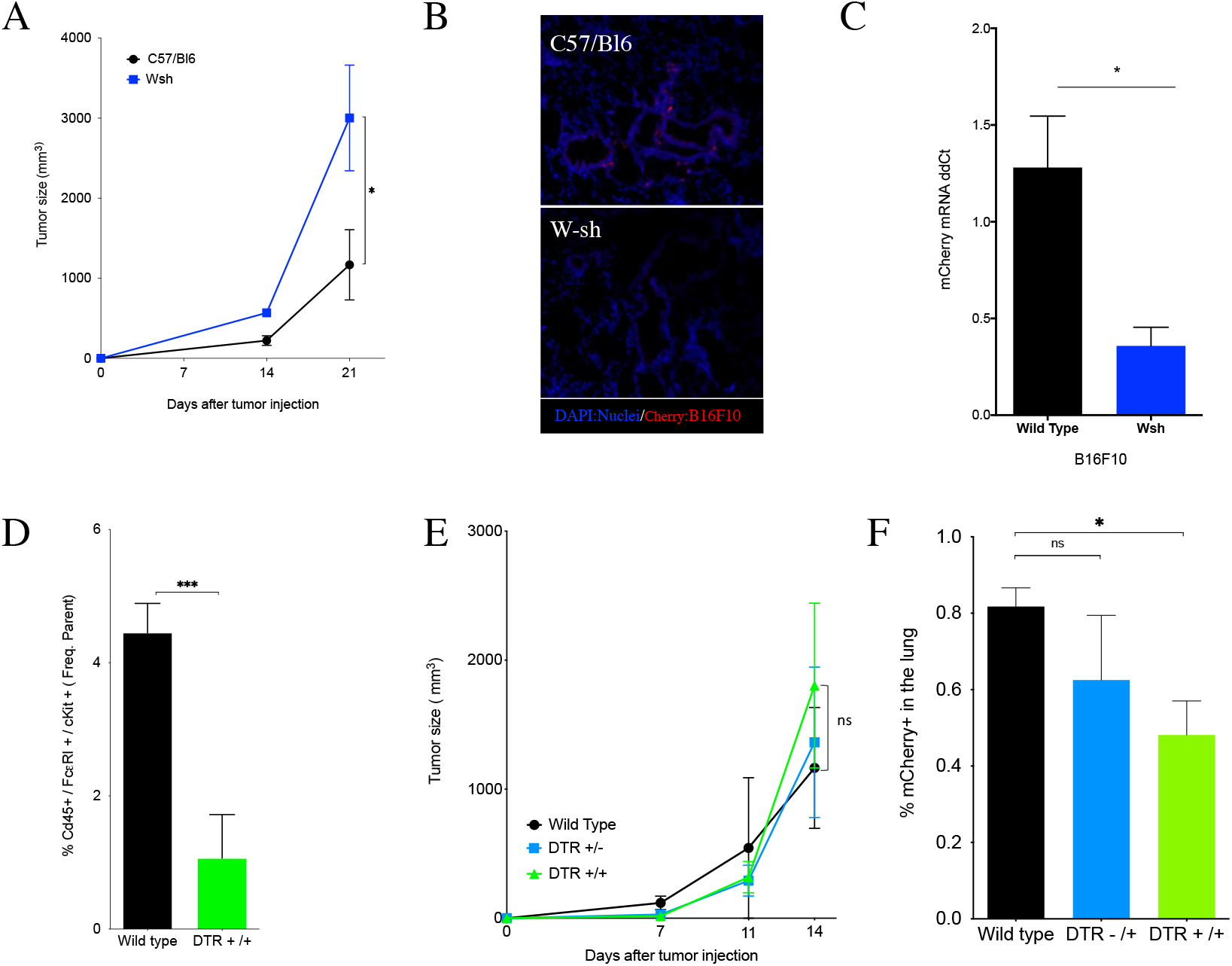
Mast cells deficiency dampens melanoma metastasis. **A)** We injected B16F10-mCherry cells (1×10^6^) subcutaneously in C57Bl6 and Wsh mice and monitor tumor growth for up to 21 days. Tumor growth mean and S.E.M. of three replicates with 3 to 5 mice per group and replicate. *p<0.05 Parametric t-test. **B)** Tumor bearing mice showed micrometastatic foci in Wsh Lungs after 21 days but not in the other groups. Lung representative IF, showing tumor cells in red (mCherry) and nuclei in blue (DAPI) staining. **C)** mCherry expression in mouse lungs after 21 days, detected by qPCR. mCherry mRNA copies in B16F10-mCherry Wsh lungs were increased when compared with C57Bl6 mice. N=3 (3 to 10 mice) p<0.05 Parametric t-test. **D)** Mast cell populations in murine *wt* and MCTP5-DTR lungs detected by flow cytometry. Phenotypic characterization: Cd45+/Fcε-RI+/cKIT+.***p<0.005 nonparametric t-test. **E)** B16F10-mCherry cells injected (1×10^6^) subcutaneously in *wt* (C57Bl6), and MCTP5-DTR +/+ (DTR+/+) mice. MCTP5-DTR +/- (DTR+/-) mice were used as littermate controls. We monitor tumor average growth twice a week. N=2, with 6 to 13 animals per group in total. **F)** B16F10 mCherry detection in *wt*, DTR +/+ and DTR+/- dissociated lungs as detected by flow cytometry. *p<0.05 Parametric t-test

### Bone-marrow derived mast cells reduce melanoma cell homing and metastasis in lungs after co-incubation *in vitro*

Our data, together with recent reports (16), support MCs contribution to melanoma metastasis. During melanoma progression MCs are influenced by tumor cells. However, there are reports showing their dual role in cancer (12, 20, 21) combining both pro- (22-24) and anti-tumor effects (25, 26) To understand the innate role of MCs, we used BMDMCs derived from *wt* mice and analyzed their influence in melanoma cell behavior. We co-incubated (CI) *in vitro* B16-F10 melanoma cells with murine BMDMCs in a 1:1 ratio for 30 minutes, injecting then the mix intravenously. Surprisingly, while B16-F10 control cells form macrometastasis in the lung 14 days after the injection, the CI of B16-F10 with BMDMCs significantly reduced lung metastasis (Figure 2A, B). To understand if CI was necessary to reduce metastatic ability of melanoma cells, we injected the same amount of melanoma cells and BMDMCs sequentially without previous CI (Fig. 2C, D). We observed that sequential injection *in vivo* did not significantly reduce lung metastasis, but co-incubation did (Fig. 2D). In order to analyze the mechanism involved, we evaluate cell death, but we did not detect any significant changes in apoptosis (Supp. Fig 2).

**Figure 2.**
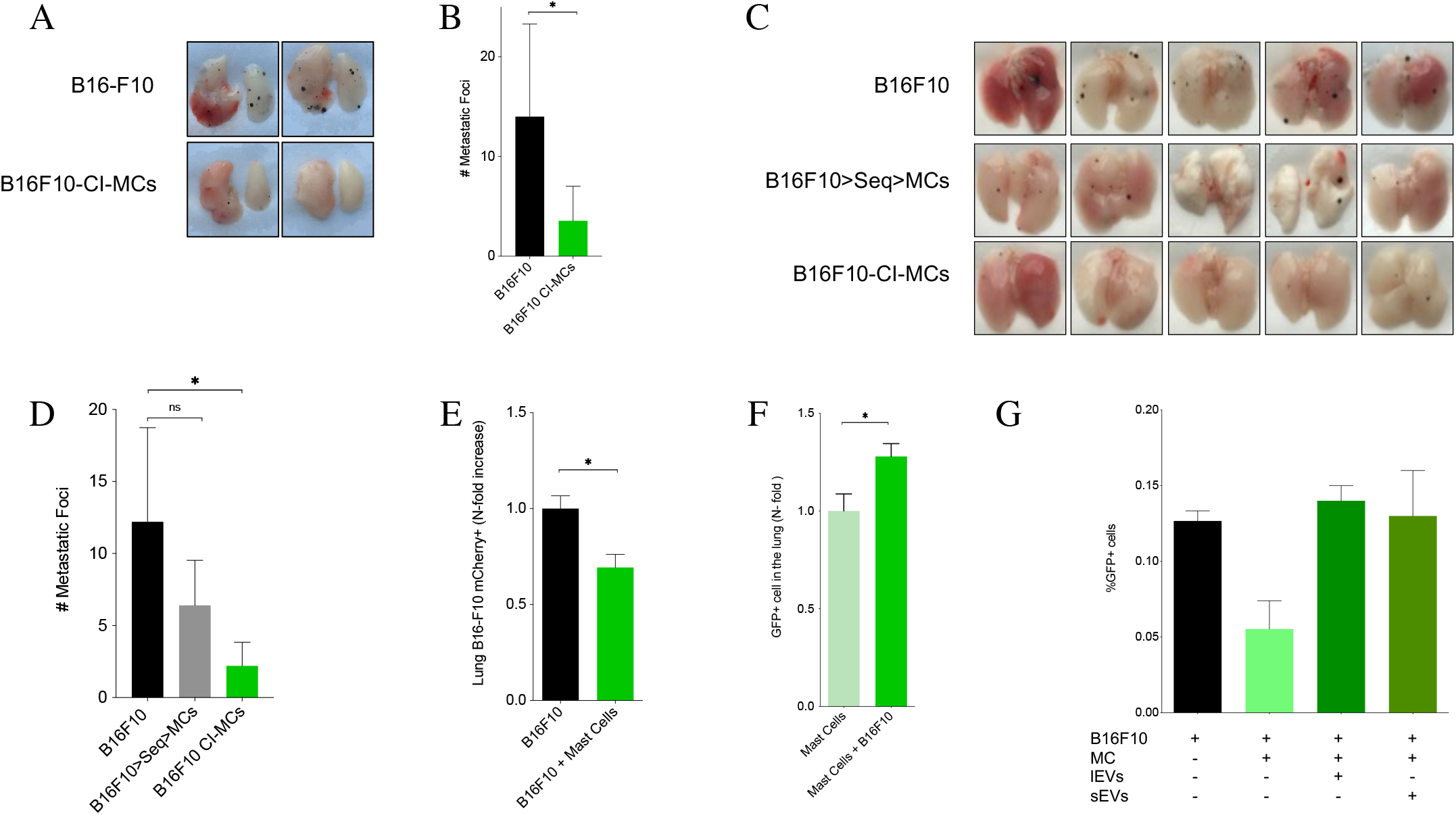
Mast cells and melanoma coincubation reduce tumor growth and metastasis to the lung. **A)** Tail vein injection of melanoma and MCs coincubation. MCs and B16F10 were coincubated (Ratio 1:1, 1×10^5^ :1×10^5^ cells) for 30 minutes and then injected in through the tail vein. Representative pictures of lungs 14 days after injection. **B)** Macro-metastatic foci quantification. N=4, 10 to 18 mice in total. * p<0.05 non-parametric t-test, **C)** B16F10 cells were injected in the tail vein of C57Bl6. another group received two sequential injections: B16F10 and then MCs cells. The third group received a coincubation of B16F10 and MCs (Ratio 1:1, 1×10^5^ :1×10^5^ cells) **D)** Metastatic foci quantification. N=5 mice. *P< 0.05 non-parametic t-test. E, Tail vein injection of melanoma and MCs coincubation after 6 hours. MCs and B16F10 cells were coincubated (Ratio 1:1, 1×10^6^ :1×10^6^ cells) for 30 minutes and then injected in C57bl6 mice. Animals were euthanized after six hours, and lungs were processed immediately for flow cytometry analysis of mCherry detection in the lung. **F)** mBMDMC-GFP Control and B16F10 cells were coincubated (Ratio 1:1, 1×10^6^ :1×10^6^ cells) for 30 minutes and then injected subcutaneously in C57bl6 mice tail vein. Animals were euthanized after six hours, and lungs were processed immediately for flow cytometry analysis of GFP+ cells in the lung. N=3, 9 mice per group in total. * p< 0.05 nonparametric t-test. **G**) MCs GFP+ detection in lung. B16F10 cells were coincubated with MCs (Ratio 1:1, 1×10^6^ :1×10^6^ cells) or treated with 10 µg of Microvesicles and EVs for 30 minutes and then injected subcutaneously in C57bl6 mice tail vein. Animals were euthanized after six hours, and lungs were processed immediately for flow cytometry analysis of mCherry detection in the lung

We next analyzed if tumor cell homing in the lungs was affected by CI. We evaluated tumor cell presence in lungs 6 hours after intravenous injection. We observed a significant reduction of tumor cell homing after CI with BMDMCs (Figure 2E, Supp. Fig 3A). We also observed an increase of GFP+ BMDMCs in the lungs in this experimental setting (Figure 2F and Supp. Fig 3B), suggesting that BMDMCs colonize the lung microenvironment reducing tumor cell homing. To test if besides physical interaction during the CI, MC-secreted factors could be involved in this effect, we investigated of the capacity of MC-derived extracellular vesicles (EVs) or soluble factors to inhibit melanoma metastasis. Co-injection of tumor cells with MC-derived large EVs (lEVs) and MC-derived small EVs (sEVs) did not affect tumor cell homing (Fig. 2G). Therefore, our data support the idea that tumor-BMDMCs co-incubation *in vitro* impairs tumor cell homing and metastasis through direct contact effect rather than secreted factors.

### HMGA-1 reduction in melanoma cells diminish metastatic behavior

To define the mechanism involved, we analyzed the secretome before and after melanoma cells-BMDMC CI. Proteomic profiling of secreted factors revealed 57 proteins significantly altered in the secretome after co-incubation of B16F10 with BMDMC (Figure 3A, 3B). Out of the many candidates HMGA1 secretion was the top protein significantly downregulated after co-incubation with BMDMCs (Fig. 3A, 3B). Since the HMG protein family is associated with cancer and tumor progression (27), we analyzed the role of HMGA1 in melanoma. First, we analyzed HMGA1 expression in a panel of mouse and human melanoma cells, we found that HMGA1 was expressed in B16-F1 and B16-F10 mouse melanoma cells (Figure 3C) as well as in the human cells analyzed (Figure 3D). Next, we confirmed the reduction of HMGA1 secretion after CI with BMDMCs in B16F1 and B16F10 (Figure 3E). Due to the reduction of metastasis observed concomitantly with HMGA1 secretion by CI, we hypothesized that HMGA1 may have a pro-metastatic role in melanoma. HMGA1 has been involved in metastasis in several types of cancer (18, 28-30), however its role in melanoma remains undefined. To get further insights, we knocked down HMGA1 expression in B16F10 (Figure 3F) and then analyzed their metastatic ability by performing experimental metastasis assays after intravenous injection. We analyzed metastasis macroscopically 14 days after injection and found that interference of HMGA1 in B16F10 (B16F10-ShHMGA1) reduced both number and size of lung metastases (Figure 3G and H). We also analyzed metastatic dissemination in the lungs staining with HMB-45, a bona fide melanoma marker (31), and found that HMGA1 downregulation significantly reduced B16F10 cells metastatic dissemination, supporting a pro-metastatic role for HMGA1 in melanoma (Figure 3I).

**Figure 3.**
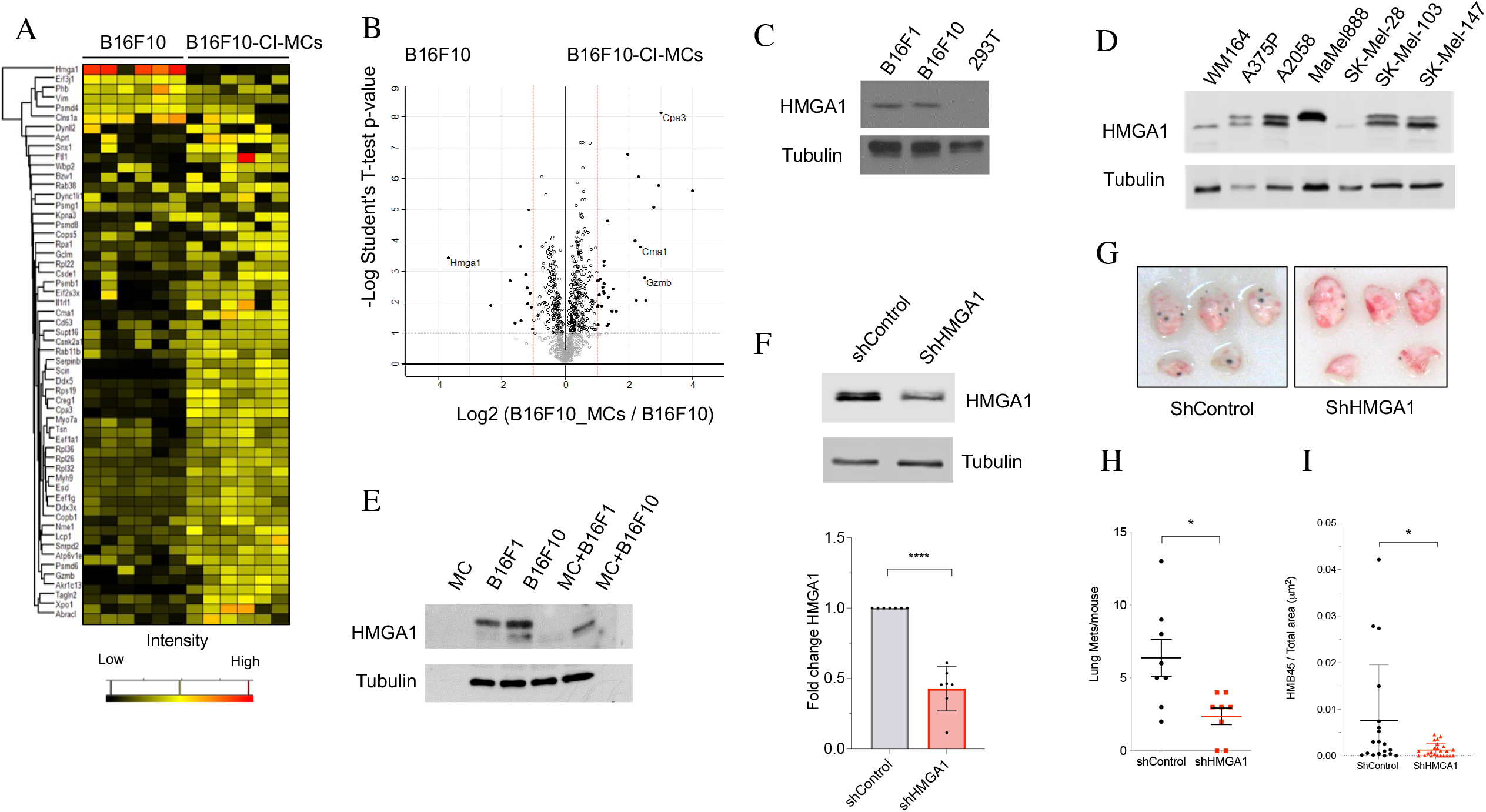
Melanoma and MC coincubation identifies HMGA1 as a melanoma metastasis driver. **A)** Proteomic analysis of melanoma secretome. B16F10 and B16F10-CI-MCs coincubation secretome was analyzed by quantitative proteomics. We show 57 significantly regulated proteins. Among them, HMGA1 is the most abundant protein in B16F10 secretome, being almost absent in the CI secretome. **B)**, Volcano plot representing the 57 significantly regulated proteins comparing B16F10 and B16F10-CI-MCs secretomes. HMGA1 is highly expressed in B16F10 secretome. Typical Mast cells proteins are present in the coincubation secretome, as Chymase (CMA1) or Mast cell Carbopeptidase (CPA3). **C, D)**. Representative Western blot for HMGA1 expression in murine (C) and human (D) melanoma cell lines. **E)** Representative Western blot of the concentrated secretome of melanoma cell lines and coincubation with MCs. MCs, B16F1, B16F10 and coincubation of MCs and B16F1 or B16F10 were resuspend in PBS, and incubated for 30 minutes at RT. Then, supernatant was concentrated and the whole reduced volume was analyzed by WB. **F)** B16F10 cells were infected with lentiviral particles to reduce HMGA1 expression (shHMGA1). A scramble sequence was used as control (shControl). B16F10 shControl and B16F10 shHMGA1 (1×10^5^) cells were injected intravenously in an experimental metastasis model, observing a decrease in metastatic foci in the B16F10sh HMGA1 injected animals. Representative image of a lung is depicted (n=8). **G)**, Representative images of tail vein injection model. B16F10 shControl and B16F10 shHMGA1 lungs, and H) lung macrometastases quantification after 14 days post injection. **I)** HMB45 staining quantification of lung tissue obtained from experimental metastasis model described in G.

### HMGA1 cytoplasmic expression in metastatic lesions correlates with reduced overall survival in melanoma patients

Human HMGA1 expression promotes tumor invasion and metastasis in several cancer types (17), however the role in melanoma is unknown. Therefore, we decided to analyze in detail the expression of HMGA1 in human melanoma. First, we analyzed HMGA1 expression in the Cancer Genome Atlas (TCGA) and we did not find significant differences in HMGA1 expression in metastatic tissue (met) when compared with primary tissues (PT) (Figure 4A). However, we hypothesized that high HMGA1 expression could influence disease outcome and analyzed the correlation of high HMGA1 expression (top 25% expression) with overall and disease-free survival. We found that high HMGA1 expression in metastatic lesions correlates with reduced overall (Figure 4B) and disease-free survival (Figure 4C). Since reduced levels of nuclear HMGA1 and increased secretion have been involved in metastatic behavior in breast cancer (18), we evaluated HMGA1 subcellular localization in 51 different primary tumors and metastatic lesions. We first confirmed that we were able to differentiate between cytoplasmic and nuclear HMGA1 localization, focusing on the cellular distribution of the signal (Supp. Figure 5) and quantified the relative abundance in these regions (Figure 4D). Analysis showed that cytoplasmic staining was increased by 4-fold in metastatic lesions compared to 1.5-fold increase showed in primary tumors (Figure 4E). These data suggest that nuclear localization of HMGA1 is reduced in metastatic lesions, concomitantly with an increase in cytoplasmic localization and potential secretion. This is consistent with the cellular distribution previously described in triple negative breast cancer, associated with metastatic disease (18).

**Figure 4.**
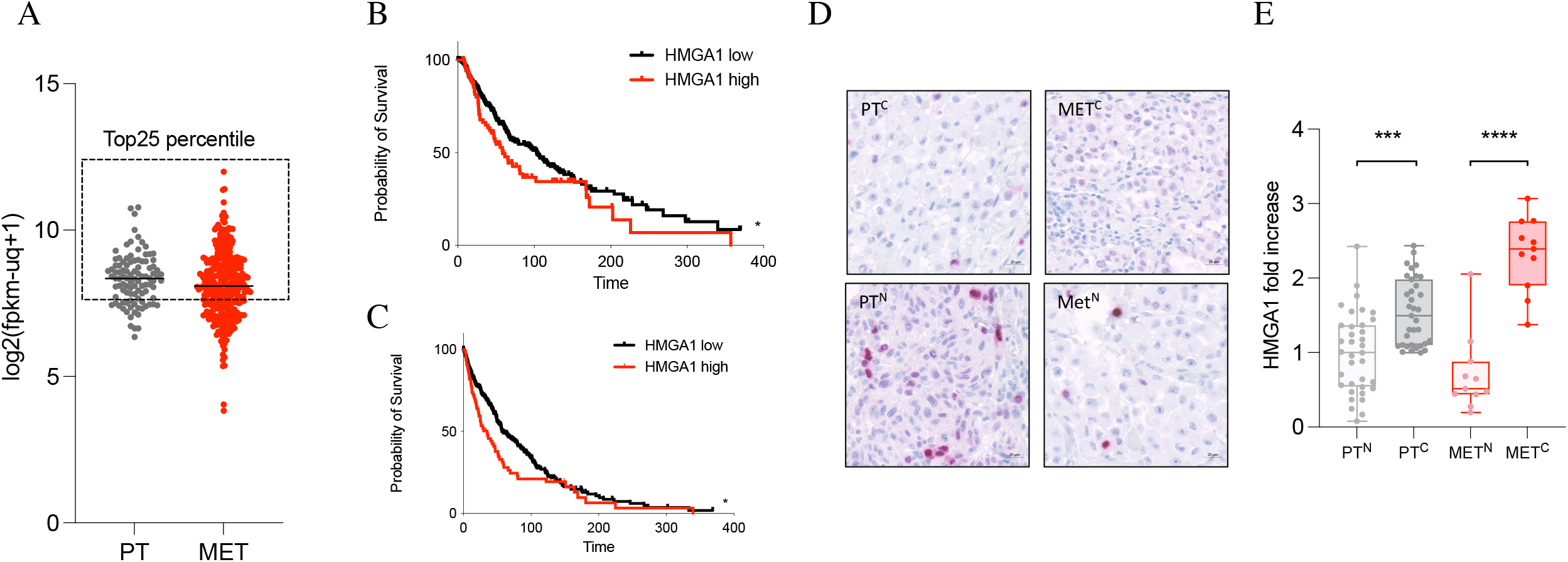
HMGA1 expression in human metastatic tissue correlated with poor prognosis. **A)** Tumor and metastatic melanoma samples with the top 25% expression levels of *HMGA1* gene (“high”) pointed with dashed square. Metastatic samples with the top 25% expression levels of *HMGA1* gene (“high”) show significantly worse prognosis than the remaining metastatic samples (“low”). **B)** Overall and **C)**, distant metastasis-free survival curves in melanoma metastatic samples expressing (top 25%) *HMGA1* from the TCGA dataset. **D)** Representative immunohistochemistry staining for HMGA-1 in human melanoma. 0= absent, 1= weak nuclear HMGA-1, 2= moderate, 3= strong. Red dashed square shows exclusive cytoplasmatic staining. **E)** Representative images of tissue microarray staining for human HMGA-1. F, Tissue Microarray quantification. Percentage of positive cells for HMGA-1 in each group. PT, Primary Tumor, MET, Metastatic tissue, C, Cytoplasmatic, N, Nuclear. Scale bar measures 20 micrometers. Kaplan-Meier survival curves were generated and differences in OS and DFS survival were assessed by log-rank Mantel-Cox test using GraphPad PRISM 8.1.1. Analyses were performed in the TCGA datasets with all tumors (101 primary tumors and 339 metastasis).

## Discussion

Melanoma metastasis is responsible for 90% of melanoma related deaths, and new approaches are urgently needed to reduce its mortality (1, 2). Recently, new findings connected tumor-derived signals with PMN formation and the role of local and recruited immune cell populations in this process (32). Mast cells, present in most barrier tissues, are innate immunity cells with a controversial role in cancer (7). Different reports have addressed the role of MCs in melanoma metastasis to the lung (10). Ablation of connective tissue mast cell (CTMC) subtype reduced melanoma metastasis in experimental models, consistent with our spontaneous metastasis results. In a different report, mice in which with 3 mast cell proteases were genetically ablated showed increased lung metastasis and reduced immune recruitment to the metastatic site (11). Therefore, it is still unclear the specific role of MCs in melanoma metastasis. Our results support a role of MCs in tumor progression, supporting the idea that MCs contribute to metastatic progression depending on the context. We investigated how MCs influence melanoma metastatic cell behavior. We observed that when melanoma cells are co-incubated with MC derived from *wt* mice in experimental metastasis models, there was a reduction of lung metastasis suggesting that co-incubation somehow reduce metastatic ability. Analysis of secreted factors after co-incubation showed that secretion of HMGA-1 by melanoma cells is reduced, suggesting a role for this factor in melanoma metastasis. HMGA1 is a chromosomal protein that binds to the minor groove of DNA and controls transcriptional activity(33). HMGA proteins are expressed at high levels during embryogenesis, but have low or undetectable levels in most adult tissues (34). HMGA1 overexpression has been correlated with metastasis and reduced patient survival in breast cancer (30, 35, 36), hepatocellular carcinoma (37) and pancreatic cancer(38). Importantly, it was recently demonstrated that HMGA1 is secreted by invasive breast cancer cells mediating cell migration, invasion, and metastasis (18). However, the role of HMGA1 in melanoma metastasis is not defined. Our data shows how co-incubation of melanoma and BMDMCs reduce melanoma metastasis and HMGA1 secretion. Reduction of HMGA1 expression in B16-F10 cells reduced melanoma metastasis suggesting a role in melanoma progression. To determine the relevance in human melanoma we analyzed HMGA1 expression in primary tumor and metastatic lesions, showing that high expression of HMGA1 in metastatic tissue is associated with decreased overall survival. Moreover, the analysis of the subcellular localization of HMGA1 in an additional cohort of melanoma primary tumors and metastasis showed a significant increase in cytoplasmatic HMGA1 expression. Importantly, increased cytoplasmic localization and secretion of HMGA1 has been previously associated with increased metastatic behavior in triple negative breast cancer (29), suggesting that a similar model might happen in melanoma. Here we present the first evidence that HMGA1 plays a role in melanoma metastatic behavior and that BMDMCs execute an anti-tumor response by inhibiting its secretion by co-incubation with tumor cells. We present evidence that reducing levels of HMGA1 in melanoma cells can indeed reduce their metastatic behavior. Therefore, therapeutic strategies targeting HMGA1 in melanoma cells could represent novel approach to reduce melanoma metastasis.

## Material and Methods

### TCGA data analysis

To determine the clinical value of HMGA1 expression in human melanoma samples, we retrieved the gene expression data and its associated clinical information of the study from the Cancer Genome Atlas Network (TCGA). GDC TCGA Melanoma (SKCM) data were extracted from cBioPortal: http://www.cbioportal.org/. The normalized expression of HMGA1 was available for 440 melanoma samples: 101 primary tumors, 339 metastases. For HMGA1 gene, normalized expression was categorized as “High expression” when it was above the third quartile (top 25% expression) of all tumor and metastatic samples; otherwise, it was categorized as “low”. Kaplan-Meier estimates of HMGA1 expression (assessed separately for each stratum) were plotted and compared to overall survival (OS), disease-free survival (DFS) curves using log-rank and Chi-square tests.

### Tumor samples

Briefly, this study was performed in two independent data samples: discovery series, which included 51 malignant melanoma, and validation series composed by 6 malignant primary tumor melanoma and their correspondent metastasis. Tissue microarray containing 53 melanoma and metastatic samples (15 of them including non-tumoral paired tissue) and associated data used in this Project were provided by the CNIO-Biobank (B.000848) after the appropriate ethic approval. This study was approved by the local ethical committee from each institution, and a complete written informed consent was obtained from all patients.

### Immunohistochemistry (IHC)

HMGA1 expression and subcellular localization was analyzed by IHC in TMAs and in PDXs, according to standard methods as previously published (18). HMGA1 monoclonal antibody was generated as previously described (18), and HMGA1 expression was classified as positive when more than 10% of tumor cells showed positive staining. Cytoplasmic and nuclear staining of HMGA1 was quantified analyzing the percentage of positive immunolabeled cells over the total cells in each tissue area Also, the overall percentage of positive cells was recorded, as well as the pathologist’s semiquantitative estimate of the overall staining intensity (0-3+). The cases were also designated as “positive” if more than 10% of the tumor cells showed expression of the antigen. All studied tumor sections also included normal tissue as an internal control. In negative controls, the primary antibodies were omitted.

### Mice

C57BL/6 females (C57), C57BL/6-Tg (CAG-EGFP)131Osb /LeySopJ (GFP mice) and B6.Cg-KitW-sh/HNihrJaeBsmJ (W-sh) of 8-12 weeks-old were purchased from the Jakson Laboratory. Wsh mice present embryonic deficit and eventual abolishment of mast cells soon after birth. Dr. Axel Roers kindly provided MCPT-5 Cre-R-DTA mice (MCTP5 DTR +/+)(19) which lack connective tissue mast cells. We used MCPT-5 Cre-R-DTA heterozygous (MCTP5 DTR +/-) mice as controls. For primary tumor growth and spontaneous lung metastasis experiments, we injected 1 × 10^6^ B16-F10 mCherry cells subcutaneously in the mice right flank. We monitored tumor growth at least thrice a week. Metastasis was evaluated by mCherry expression in the lung after two or three weeks. For education experiments, we injected a total of 10µg of EVs every 72h for 2 or 3 weeks in the eye sinus. Synthetic unilamellar 100 nm liposomes (Encapsula Nanoscience) or PBS were used as controls in all studies. For lung colonization experiments, 1 × 10^6^ B16-F10–mCherry cells were injected, and C57BL/6 female mice were euthanized after 6 hours. For experimental metastasis, we injected 1 × 10^5^ B16-F10– mCherry cells, and C57BL/6 female mice were euthanized after two weeks. Animals were sacrificed and tumor and lung samples were dissected and frozen at -80C in O.C.T. All mouse work was performed in accordance with institutional, IACUC and AAALAS guidelines. All animals were monitored for abnormal tissue growth or ill effects according to AAALAS guidelines and we euthanized animals if excessive health deterioration was observed.

### Melanoma Cell lines

B16F10 and B16F1 were purchased from ATCC. B16F10-mCherry cells were generated as previously described(39). All melanoma cells were grown in DMEM supplemented with 10% (v/v) sEV-reduced fetal bovine serum (FBS) glutamine 2mM and antibiotics. Human melanoma cells were kindly provided by Marisol Soengas.

### Tissue processing

Specimens were fixed in 4% PFA. For immunofluorescence, 12 μm O.C.T. tissue sections of lungs were treated with PBS-0.3% triton X-100 for 15 min. Non-specific sites were blocked by incubation in PBS containing PBS 1% BSA 5% and specific Serum – 0.05% triton X-100 for 1h at RT. We incubated primary antibody, anti-mCherry (ab167453, Abcam), overnight at 4C. After washes with PBS, lungs were incubated 1 h with secondary antibodies from Alexa Fluor series from Molecular Probes (dilution 1:500) and washed again. Finally, samples were mounted with Prolong-DAPI(Thermo). Fluorescent images were obtained using a Nikon confocal microscope (Eclipse TE2000U) and analyzed using Nikon software (EZ-C1 3.6).

### Flow Cytometry analysis

For lung population analysis, tissues were minced and then digested at 37°C for 20 min with an enzyme cocktail (collagenase A, dispase and DNaseI, Roche Applied Science). Single-cell suspensions were prepared by filtering through a 70-μm strainer and passing through a nylon mesh. We eliminate red blood cells with ACK lysis buffer (Gibco), and then cell suspensions were washed in Flow buffer (PBS, 5mM EDTA and 1% BSA) and incubated with the following primary antibodies: anti-mouse CD45 (Brilliant Violet 570, 30-F11, 1:200, Biolegend), anti-mouse Fce-R1 alpha (PE-Cyanine7, MAR-1, eBioscience™ 1:400), anti-mouse cKIT (APC/Cy7, 2B8, 1:200, Biolegend). To assess cell viability, we used LIVE/DEAD™ Fixable Violet Dead Cell Stain Kit, for fixed cells (Thermo, L34963) and Pacific Blue™ Annexin V Apoptosis Detection Kit with 7-AAD, (Biolegend, 640926). For in vitro experiments, adherent cells were treated with cell dissociation buffer (Gibco) and single cell suspensions were washed in Flow buffer prior to primary antibody incubation. GFP+ & mCherry+ presence in the lung were analyzed using a FACSCalibur or a FACSCanto (Beckton Dickinson). FACS data was analyzed with FlowJo software (TreeStar Inc.).

### qPCR

We isolated RNA from frozen lungs using Qiagen RNeasy Mini Kit, to then perform reverse transcription (SuperScript™ VILO™ cDNA Synthesis Kit, Thermo) and quantitative PCR for specific gene expression using pre-designed TaqMan® assays (Thermo). GFP and mCherry expression was assessed using specific primers(39) and SybrGreen PCR reagents (Applied Biosystems). Quantitative real-time PCR (QRT-PCR) was performed on a 7500 Fast Real Time PCR System (Applied Biosystems). Relative expression was calculated following delta delta Ct calculation method and *Beta-Actin* and *18s* were used as housekeeping genes.

### Ex vivo Mast cell differentiation

Bone Marrow derived mast cells were obtained after flushing tibia and femur from C57Bl6 (mBMDMCs) and C57BL/6-Tg (CAG-EGFP)131Osb /LeySopJ (mBMDMCs-GFP) as previously described. Briefly, bone marrow progenitors were cultured in OPTIMEM supplemented with sEV-reduced FBS and 5ng/ml of interleukin-3 for 4 weeks. We maintained cell density at 6×10^5^ cells/ml and evaluate cKIT and FcE-R1 expression weekly to validate differentiation.

### Extracellular vesicle (EV) isolation

We performed small EV isolation by sequential Ultracentrifugation. Cells were cultured in media supplemented with 10% sEV-reduced FBS (FBS, Hyclone). FBS was reduced of bovine sEVs by ultracentrifugation at 100,000xg for 70 min. Supernatant fractions collected from 72-96 h cell cultures were pelleted by centrifugation at 300xg for 5 min and 500xg for 10 min. The supernatant was centrifuged at 12,000xg for 20 min, and the pellet collected for large EV analysis. sEVs were then harvested by centrifugation of previous step supernatant at 100,000xg for 70 min. The sEV pellet was resuspended in 20 ml of PBS and collected by ultracentrifugation at 100,000g for 70 min. All spins were performed at 10ºC in Beckman instruments. When indicated, EVs were labeled with PKH26 (Sigma). Labeled EVs were washed twice in 20 ml of PBS, collected by ultracentrifugation and resuspended in PBS using standard techniques as described above.

### Coincubation experiments and secretome analysis

Same number of mBMDMCs and B16F10 cells (1×10^6^) were pellet down and washed with PBS. After washing, cell pellets were resuspended in 500 ul of PBS and mixed for coincubation for 30 minutes at RT. The conditioned media were centrifugated 300xg, 5 min; 500xg, 10 minutes and 3000xg, 20 minutes. Conditioned media was frozen for proteomic analysis or was concentrated using a 10,000 MWCO Millipore Amicon Ultra (Millipore) to further evaluate secretome protein expression.

### Western blot

Whole cell lysates, microparticles, EVs and conditioned media were resolved by SDS-PAGE and Western blot analysis was performed with the following primary antibodies: rabbit anti-HMGA1 (sc- 393213, Santa Cruz Biotechnology), mouse anti-alpha-tubulin (X), β-actin (Sigma Aldrich, #A5441). Mouse m-IgGκ BP-HRP (sc-516102, Santa Cruz Biotechnology) and Peroxidase conjugated AffinityPure donkey anti-Rabbit or anti-Mouse (1:5000, Jackson ImmunoResearch, #711-035-152 and 1:5000, #715-035-150, respectively) were used as secondary antibodies. ECL Western Blotting Substrate (GE Healthcare) Kit was used for western blot developing. The intensities of the immunoreactive bands were quantified by densitometry using ImageJ software (NIH).

### Proteomic analysis of the secretome

Samples were lysed in urea and digested with Lys-C/trypsin using the standard FASP protocol. Peptides were analysed by LC-MS/MS analysis using an LTQ Orbitrap Velos mass spectrometer (Thermo Scientific). Raw files were analysed with MaxQuant against a mouse protein database and the MaxLFQ algorithm was used for label-free protein quantification. The MS proteomic data have been deposited to the ProteomeXchange Consortium via the PRIDE partner repository with the dataset identifier PXD009505. For peer reviewing purposes, the data set can be available under the username and password: RySIl4Q1.

### Lentiviral HMGA1 knock-down

To stably knockdown HMG-I/HMG-Y we used lentiviral particles (Santa Cruz Biotechnology) either containing 3 target-specific shRNA targeting mouse HMG-I/HMG-Y (sc-37116-V) or control shRNA lentiviral particles targeting a scramble sequence (sc-108080). B16F10 cells were infected overnight with lentiviral particles in the presence of polybrene (Sigma, TR-1003). Stable cell lines were obtained after one-week selection with puromycin and HMGA1 depletion was confirmed by immunoblot. Antibodies: HMGA1 (Santa Cruz biotechnology, sc-393213; 1:1000), or β-ACTIN (Cell Signaling, 4967; 1:5000) overnight at 4ºC. For HMGA1, m-IgGκ BP-HRP (Santa Cruz Biotechnology) was used as secondary antibody. For quantification, the ratio between the HMGA1 and β-ACTIN band intensities for each sample was measured using ImageJ software.

### Statistical analysis

Error bars in graphical data represent means ± s.e.m. Mouse experiments were performed at least triplicate, using at least 5 mice per treatment group, except noted. All *in vitro* experiments were performed at least in duplicate. Statistical significance was determined using Parametric & Non-parametric t-tests. p-values of *P* < 0.05 were considered statistically significant using GraphPad Prism software.

## Acknowledgments

We would like to thank all the members of the Peinado and Lyden laboratories for their insights. We would like to thank Axel Roers for the MCT5Cre mice donation and Marisol Soengas for providing human metastatic models.

## Authors contribution

A.B-M. designed and supervised the study, interpreted, and analyzed the data, and wrote the manuscript. LN. performed experiments, analyzed, interpreted, and discussed the data. M.H performed human samples experiments, managed sample cohorts, interpreted the data. E.C., E.G., M.C. and P.S-C. performed experiments, analyzed the data, and contributed to data discussions. W.B. managed colony breeding and supported the animal work. P.X-E and J.M. performed proteomic and bioinformatic analysis. I.M. discussed the experimental design and data and edited the manuscript. J.V provided the human HMGA1 antibody and discussed the results. H.P. directed and supervised the work.

## Conflict of interest

Authors declare no conflict of interest.

## Notes

### Competing Interest Statement

The authors have declared no competing interest.

## References

1. Garbe C, Amaral T, Peris K, Hauschild A, Arenberger P, Bastholt L, et al. European consensus-based interdisciplinary guideline for melanoma. Part 1: Diagnostics - Update 2019. Eur J Cancer. 2020;126:141–58.

2. Garbe C, Amaral T, Peris K, Hauschild A, Arenberger P, Bastholt L, et al. European consensus-based interdisciplinary guideline for melanoma. Part 2: Treatment - Update 2019. Eur J Cancer. 2020;126:159–77.

3. Peinado H, Zhang H, Matei IR, Costa-Silva B, Hoshino A, Rodrigues G, et al. Pre-metastatic niches: organ-specific homes for metastases. Nat Rev Cancer. 2017;17(5):302–17.

4. Galli SJ, Borregaard N, Wynn TA. Phenotypic and functional plasticity of cells of innate immunity: macrophages, mast cells and neutrophils. Nat Immunol. 2011;12(11):1035–44.

5. Bulfone-Paus S, Nilsson G, Draber P, Blank U, Levi-Schaffer F. Positive and Negative Signals in Mast Cell Activation. Trends Immunol. 2017;38(9):657–67.

6. Wernersson S, Pejler G. Mast cell secretory granules: armed for battle. Nat Rev Immunol. 2014;14(7):478–94.

7. Dimitriadou V, Koutsilieris M. Mast cell-tumor cell interactions: for or against tumour growth and metastasis? Anticancer Res. 1997;17(3A):1541–9.

8. Gurish MF, Austen KF. Developmental origin and functional specialization of mast cell subsets. Immunity. 2012;37(1):25–33.

9. Stone KD, Prussin C, Metcalfe DD. IgE, mast cells, basophils, and eosinophils. J Allergy Clin Immunol. 2010;125(2 Suppl 2):S73–80.

10. Ohrvik H, Grujic M, Waern I, Gustafson AM, Ernst N, Roers A, et al. Mast cells promote melanoma colonization of lungs. Oncotarget. 2016;7(42):68990–9001.

11. Grujic M, Paivandy A, Gustafson AM, Thomsen AR, Ohrvik H, Pejler G. The combined action of mast cell chymase, tryptase and carboxypeptidase A3 protects against melanoma colonization of the lung. Oncotarget. 2017;8(15):25066–79.

12. Derakhshani A, Vahidian F, Alihasanzadeh M, Mokhtarzadeh A, Lotfi Nezhad P, Baradaran B. Mast cells: A double-edged sword in cancer. Immunol Lett. 2019;209:28–35.

13. Pittoni P, Colombo MP. The dark side of mast cell-targeted therapy in prostate cancer. Cancer Res. 2012;72(4):831–5.

14. Ch’ng S, Sullivan M, Yuan L, Davis P, Tan ST. Mast cells dysregulate apoptotic and cell cycle genes in mucosal squamous cell carcinoma. Cancer Cell Int. 2006;6:28.

15. Ch’ng S, Wallis RA, Yuan L, Davis PF, Tan ST. Mast cells and cutaneous malignancies. Mod Pathol. 2006;19(1):149–59.

16. Ghouse SM, Polikarpova A, Muhandes L, Dudeck J, Tantcheva-Poor I, Hartmann K, et al. Although Abundant in Tumor Tissue, Mast Cells Have No Effect on Immunological Micro-milieu or Growth of HPV-Induced or Transplanted Tumors. Cell Rep. 2018;22(1):27–35.

17. Wang Y, Hu L, Zheng Y, Guo L. HMGA1 in cancer: Cancer classification by location. J Cell Mol Med. 2019;23(4):2293–302.

18. Mendez O, Peg V, Salvans C, Pujals M, Fernandez Y, Abasolo I, et al. Extracellular HMGA1 Promotes Tumor Invasion and Metastasis in Triple-Negative Breast Cancer. Clin Cancer Res. 2018;24(24):6367–82.

19. Dudeck A, Dudeck J, Scholten J, Petzold A, Surianarayanan S, Kohler A, et al. Mast cells are key promoters of contact allergy that mediate the adjuvant effects of haptens. Immunity. 2011;34(6):973–84.

20. Varricchi G, Galdiero MR, Marone G, Granata F, Borriello F, Marone G. Controversial role of mast cells in skin cancers. Exp Dermatol. 2017;26(1):11–7.

21. Biswas A, Richards JE, Massaro J, Mahalingam M. Mast cells in cutaneous tumors: innocent bystander or maestro conductor? Int J Dermatol. 2014;53(7):806–11.

22. Ribatti D, Ennas MG, Vacca A, Ferreli F, Nico B, Orru S, et al. Tumor vascularity and tryptase-positive mast cells correlate with a poor prognosis in melanoma. Eur J Clin Invest. 2003;33(5):420–5.

23. Ribatti D, Vacca A, Ria R, Marzullo A, Nico B, Filotico R, et al. Neovascularisation, expression of fibroblast growth factor-2, and mast cells with tryptase activity increase simultaneously with pathological progression in human malignant melanoma. Eur J Cancer. 2003;39(5):666–74.

24. Toth-Jakatics R, Jimi S, Takebayashi S, Kawamoto N. Cutaneous malignant melanoma: correlation between neovascularization and peritumor accumulation of mast cells overexpressing vascular endothelial growth factor. Hum Pathol. 2000;31(8):955–60.

25. Rajabi P, Bagheri A, Hani M. Intratumoral and Peritumoral Mast Cells in Malignant Melanoma: An Immunohistochemical Study. Adv Biomed Res. 2017;6:39.

26. Siiskonen H, Poukka M, Bykachev A, Tyynela-Korhonen K, Sironen R, Pasonen-Seppanen S, et al. Low numbers of tryptase+ and chymase+ mast cells associated with reduced survival and advanced tumor stage in melanoma. Melanoma Res. 2015;25(6):479–85.

27. Fusco A, Fedele M. Roles of HMGA proteins in cancer. Nat Rev Cancer. 2007;7(12):899–910.

28. Yang Q, Wang Y, Li M, Wang Z, Zhang J, Dai W, et al. HMGA1 promotes gastric cancer growth and metastasis by transactivating SUZ12 and CCDC43 expression. Aging (Albany NY). 2021;13(12):16043–61.

29. Yang M, Guo Y, Liu X, Liu N. HMGA1 Promotes Hepatic Metastasis of Colorectal Cancer by Inducing Expression of Glucose Transporter 3 (GLUT3). Med Sci Monit. 2020;26:e924975.

30. Qi C, Cao J, Li M, Liang C, He Y, Li Y, et al. HMGA1 Overexpression is Associated With the Malignant Status and Progression of Breast Cancer. Anat Rec (Hoboken). 2018;301(6):1061–7.

31. Ohsie SJ, Sarantopoulos GP, Cochran AJ, Binder SW. Immunohistochemical characteristics of melanoma. J Cutan Pathol. 2008;35(5):433–44.

32. Peinado H, Zhang HY, Matei IR, Costa-Silva B, Hoshino A, Rodrigues G, et al. Pre-metastatic niches: organ-specific homes for metastases. Nature Reviews Cancer. 2017;17(5):302–17.

33. Reeves R, Nissen MS. Cell cycle regulation and functions of HMG-I(Y). Prog Cell Cycle Res. 1995;1:339–49.

34. Chiappetta G, Avantaggiato V, Visconti R, Fedele M, Battista S, Trapasso F, et al. High level expression of the HMGI (Y) gene during embryonic development. Oncogene. 1996;13(11):2439–46.

35. Huang R, Huang D, Dai W, Yang F. Overexpression of HMGA1 correlates with the malignant status and prognosis of breast cancer. Mol Cell Biochem. 2015;404(1-2):251-7.

36. Mendez O, Perez J, Soberino J, Racca F, Cortes J, Villanueva J. Clinical Implications of Extracellular HMGA1 in Breast Cancer. Int J Mol Sci. 2019;20(23).

37. Andreozzi M, Quintavalle C, Benz D, Quagliata L, Matter M, Calabrese D, et al. HMGA1 Expression in Human Hepatocellular Carcinoma Correlates with Poor Prognosis and Promotes Tumor Growth and Migration in in vitro Models. Neoplasia. 2016;18(12):724–31.

38. Liau SS, Jazag A, Whang EE. HMGA1 is a determinant of cellular invasiveness and in vivo metastatic potential in pancreatic adenocarcinoma. Cancer Res. 2006;66(24):11613–22.

39. Peinado H, Aleckovic M, Lavotshkin S, Matei I, Costa-Silva B, Moreno-Bueno G, et al. Melanoma exosomes educate bone marrow progenitor cells toward a pro-metastatic phenotype through MET. Nat Med. 2012;18(6):883–91.

