## Supplemental figures and legend for "Mast cells impair melanoma cell homing and metastasis by inhibiting HMGA1 secretion"

### Supplementary Figures Legend

**Supplementary Figure 1.**

**Lung Mast Cells flow cytometry characterization.**

Representative flow cytometry panels of Mast cell populations in MCTP5-DTR lungs. Phenotypic characterization: SSC-W/SSC-H/Live-dead dye-/Cd45+/Fcε-RI+/cKIT+

**Supplementary figure 2**

**Mast cell- B16F10 melanoma coincubation do not induce cell death.**

**A)** Representative flow cytometry panel depicting B16F10-mCherry cells, MCs-GFP, and coincubation. **B)** Cell death Flow cytometry analysis. After 30 minutes of coincubation, coculture was stained with Annexin V and 7AAD. Cells were gated following strategy showed in A, and then necrosis and late and early apoptosis were measured. N>3.

**Supplementary figure 3**

**A)** Representative flow cytometry panels of mCherry detection in the Tail vein injected-mice lungs. **B)** Representative flow cytometry panels of GFP+ detection in the Tail vein injected-mice lungs.

**Supplementary Figure 4**

**A)** Overall survival in primary tumors and B, Distant-free survival curves in primary tumor melanoma primary tumors samples expressing (top 25%) *HMGA1* from the TCGA dataset.

**Supplementary Figure 5**

Representative image of HMGA1 staining, presenting quantification criteria: 0= absent; 1= weak nuclear HMGA-1; 2= moderate; 3= strong.

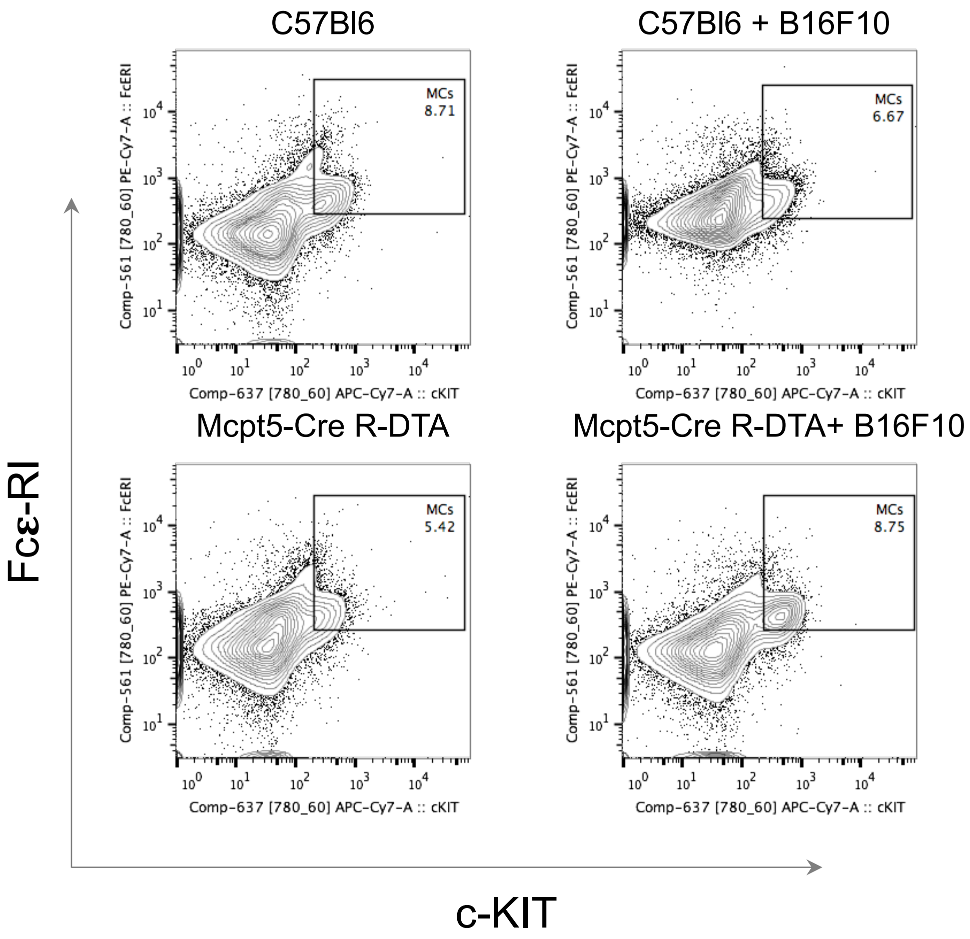


**Supplementary Figure 1**

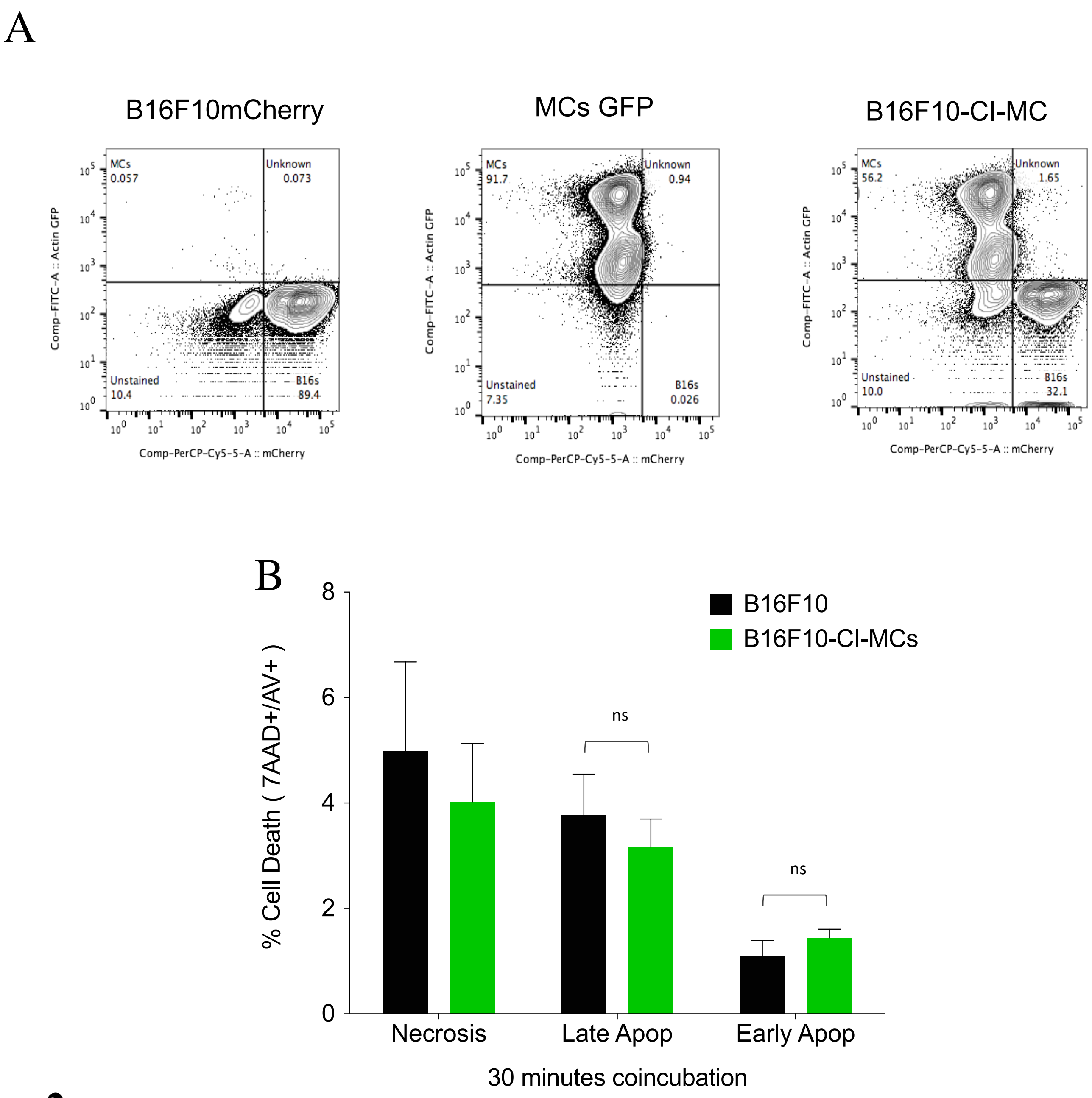


**Supplementary Figure 2**

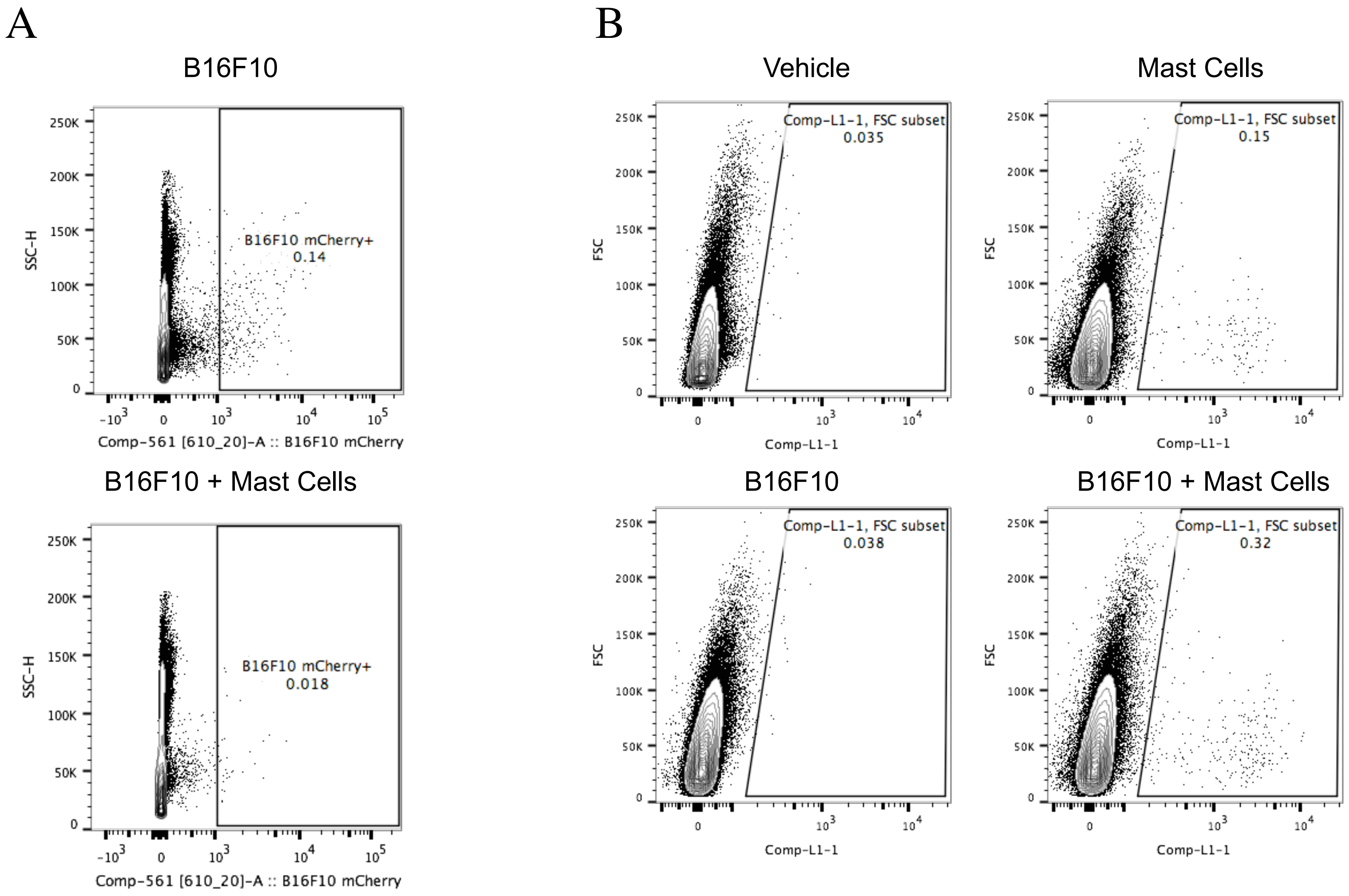


**Supplementary Figure 3**

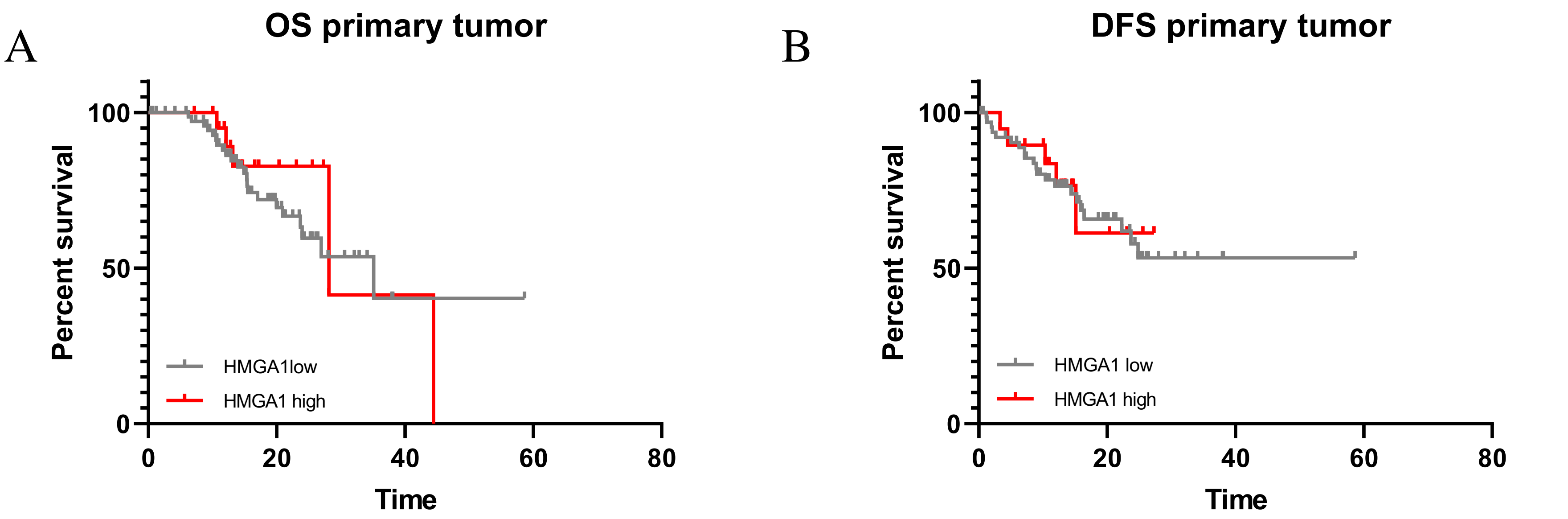


**Supplementary Figure 4**

A

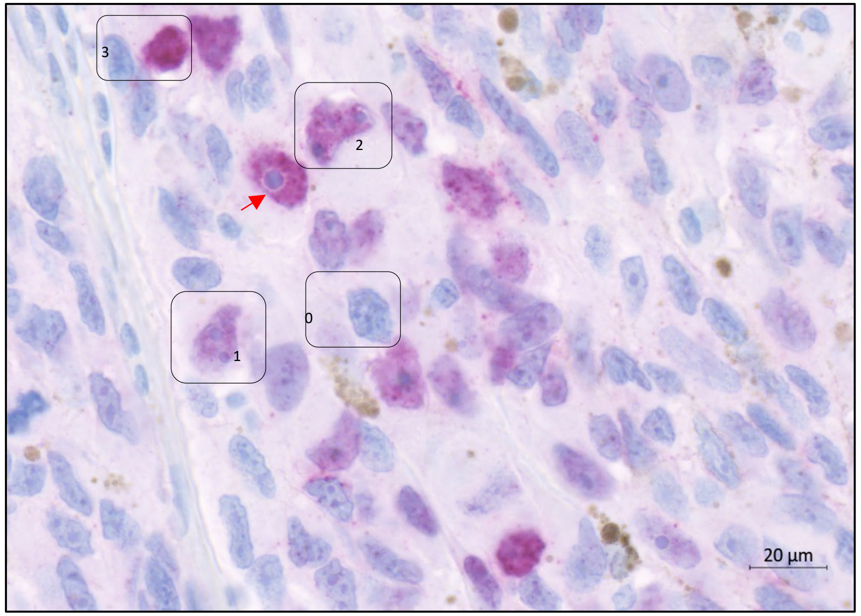


0= absent
1= weak nuclear HMGA-1
2= moderate
3= strong

**Supplementary Figure 5**
